# Single-cell selectivity and functional architecture of human lateral occipital complex

**DOI:** 10.1101/631929

**Authors:** Thomas Decramer, Elsie Premereur, Mats Uytterhoeven, Wim Van Paesschen, Johannes van Loon, Peter Janssen, Tom Theys

## Abstract

The human lateral occipital complex (LOC) is more strongly activated by images of objects compared to scrambled controls, but detailed information at the neuronal level is currently lacking. We recorded with microelectrode arrays in the LOC of two patients, and obtained highly selective single-unit, multi-unit and high-gamma responses to images of objects. Contrary to predictions derived from functional imaging studies, all neuronal properties indicated that the subsector of LOC we recorded from occupies an unexpectedly high position in the hierarchy of visual areas. Notably, the response latencies of LOC neurons were long, the shape selectivity was spatially clustered, LOC receptive fields were large and bilateral, and a number of LOC neurons exhibited 3D-structure selectivity (a preference for convex or concave stimuli), which are all properties typical of end-stage ventral stream areas. Thus, our results challenge prevailing ideas about the position of the LOC in the hierarchy of visual areas.

## Introduction

Our understanding of the human brain is hampered by the limitations imposed upon neuroscience research in humans. Noninvasive measurements of brain activity (EEG, functional Magnetic Resonance Imaging or fMRI) often provide only coarse information regarding neural activity, due to their limited spatial or temporal resolution. Genuine insight into the function of a brain area requires detailed measurements of the electrical activity of individual neurons and small populations of neurons at high spatiotemporal resolution. Intracortical electrophysiological recordings in humans are scarce, therefore the human visual cortex is virtually unexplored at the level of the individual neurons and small populations of neurons. Several studies have recorded field potentials with intracranial electrodes (Allison et al., 1999, Arroyo et al., 1993, Yoshor et al., 2007), but macroelectrode recordings still reflect activity of hundreds of thousands of neurons and, due to their large contact area, cannot measure spiking activity, nor can they reveal the microarchitecture of visual cortex on a submillimeter scale. A series of studies using depth electrodes in the mesial temporal lobe have investigated the visual responses of single neurons in entorhinal and perirhinal cortex (Fried et al., 1997, Kreiman et al., 2000b, Kreiman et al., 2000a, Kreiman et al., 2002, Quiroga et al., 2005, Quian Quiroga et al., 2009), and one study (Aflalo et al., 2015) reported single-unit responses during imagined actions (reaching and grasping) in a patient with a microelectrode array implanted in parietal cortex, who thereby obtained accurate control over a robot arm. (Self et al., 2016) measured multi-unit activity (MUA) and local field potential (LFP) activity in early visual areas (V2/V3) in a patient using hybrid macro-micro depth electrodes. This study observed that the properties of populations of neurons (multi-unit receptive fields, tuning for contrast, orientation, spatial frequency and modulation by context and attention) were similar to those of neurons in the macaque areas V2 and V3. To our knowledge, intracortical recordings in intermediate human visual areas such as the Lateral Occipital Complex (LOC) have never been performed.

Recordings in patients, combined with similar measurements in monkeys, may allow us to answer very specific questions with regard to the properties of individual neurons and the homologies between cortical areas in humans and monkeys. For example, assessing the shape selectivity or the receptive field profile of LOC neurons requires intracortical recordings, which have never been performed. Moreover, despite two decades of functional imaging studies in both species (Vanduffel et al., 2014), the homologies between the (subsectors of) LOC and ventral occipitotemporal cortex in humans and the monkey inferior temporal cortex (ITC) areas has not yet been resolved. The LO1 and LO2 subsector may be retinotopically organized (Wandell et al., 2007, Silson et al., 2013), similar to the monkey area TEO (Boussaoud et al., 1991), but direct single-cell evidence in the two species is lacking. Furthermore, the microarchitecture of these areas in humans (i.e. the spatial clustering of shape selectivity on the scale of cortical columns measuring 0.5 mm) is very difficult to assess with fMRI (Goncalves et al., 2015). To investigate the clustering of neuronal selectivity in human visual cortex, recordings with intracortical microelectrodes are necessary.

## Results

In two patients who were evaluated for refractory epilepsy, we ran an LOC-localizer fMRI experiment, in which blocks of non-scrambled shapes and outlines were interleaved with control blocks of scrambled stimuli (Fig. 1, (Kourtzi and Kanwisher, 2000)). A 96-channel Utah microelectrode array was implanted in LOC (Fig. 1A; MNI coordinates 55, −71, 1 for patient 1, and −55, −77, 6 for patient 2) (Silson et al., 2013). We verified the anatomical location of the array using a computed tomography (CT) scan obtained after array implantation, which was co-registered onto the anatomical MRI. Figure 1B shows the fMRI activations in the two patients for the contrast non-scrambled vs. scrambled stimuli, plotted on the patient’s own anatomical MRI (p<0.05, FWE corrected). The fMRI results confirmed that the microelectrode arrays were indeed implanted in the hotspot of fMRI activations.

**Figure 1.**
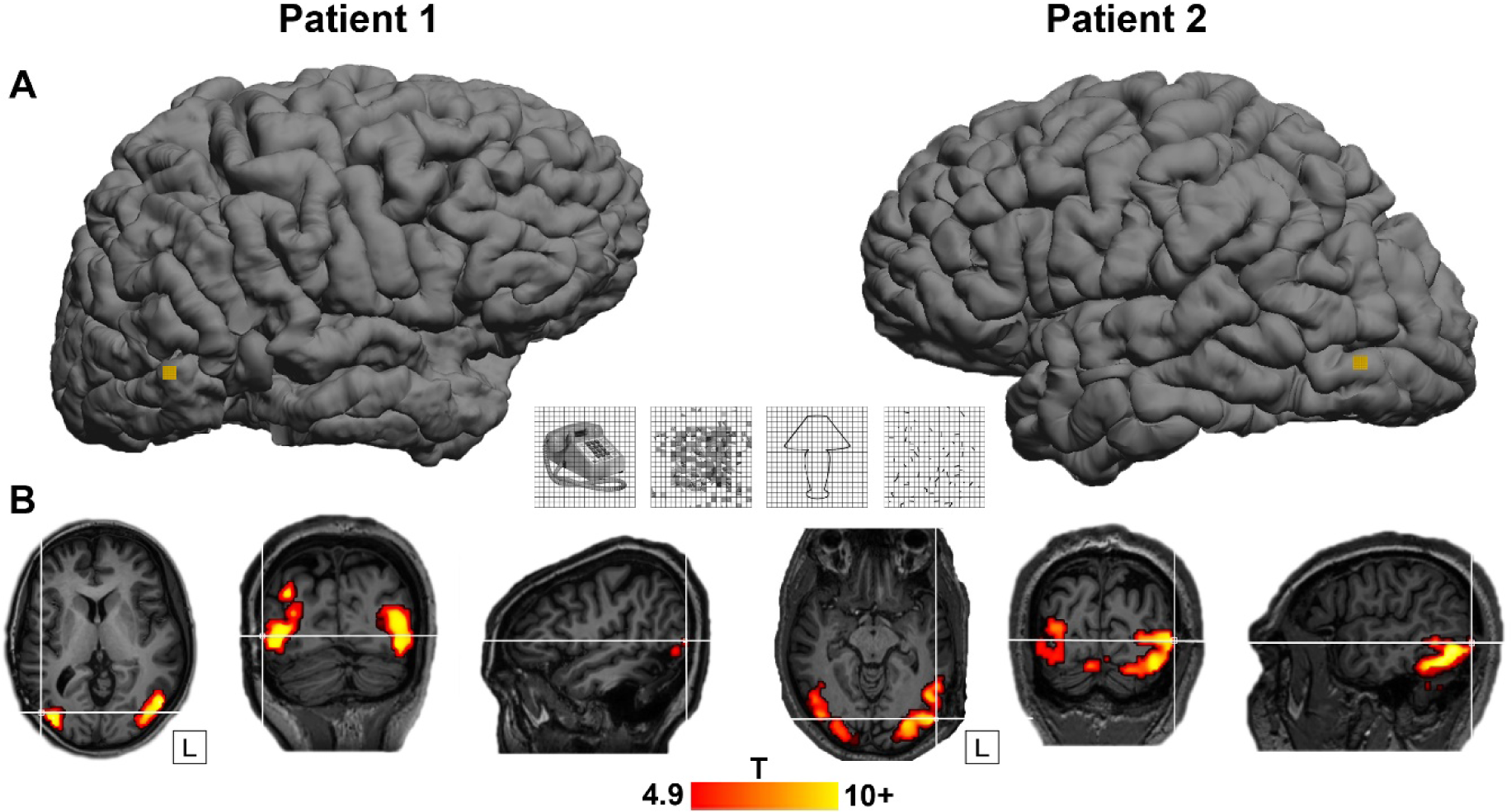
LO Localizer and location of arrays. A. Site of Utah array implantation (yellow) projected onto 3D rendering of the brain in patient 1 (left) and patient 2 (right). Center: LO Localizer stimuli (shapes, scrambled shapes, outlines, scrambled outlines). B. fMRI activations with LO localizer stimuli. T-values for contrast [shapes+outlines]-[scrambled shapes+scrambled outlines], plotted on T1 weighted image. P<0.05, FWE-corrected for multiple comparisons. Crosshair indicates the position of the Utah array in relation to the fMRI activation for both patients.

### Effect of image scrambling

The example neuron in Figure 2A responded significantly more strongly to images of objects (both shapes and outlines) compared to scrambled controls (permutation test: p <0.001, d’ = 0.72 for shapes and p = 0.002, d’ = 0.50 for line stimuli). Unlike previous recordings in the human medial temporal lobe (Kreiman et al., 2000b, Quiroga et al., 2005), this neuronal response was brisk and relatively transient (response and selectivity latency: 125 ms). The large waveform (inset in Figure 2A) indicates that this neuron was well-isolated.

**Figure 2.**
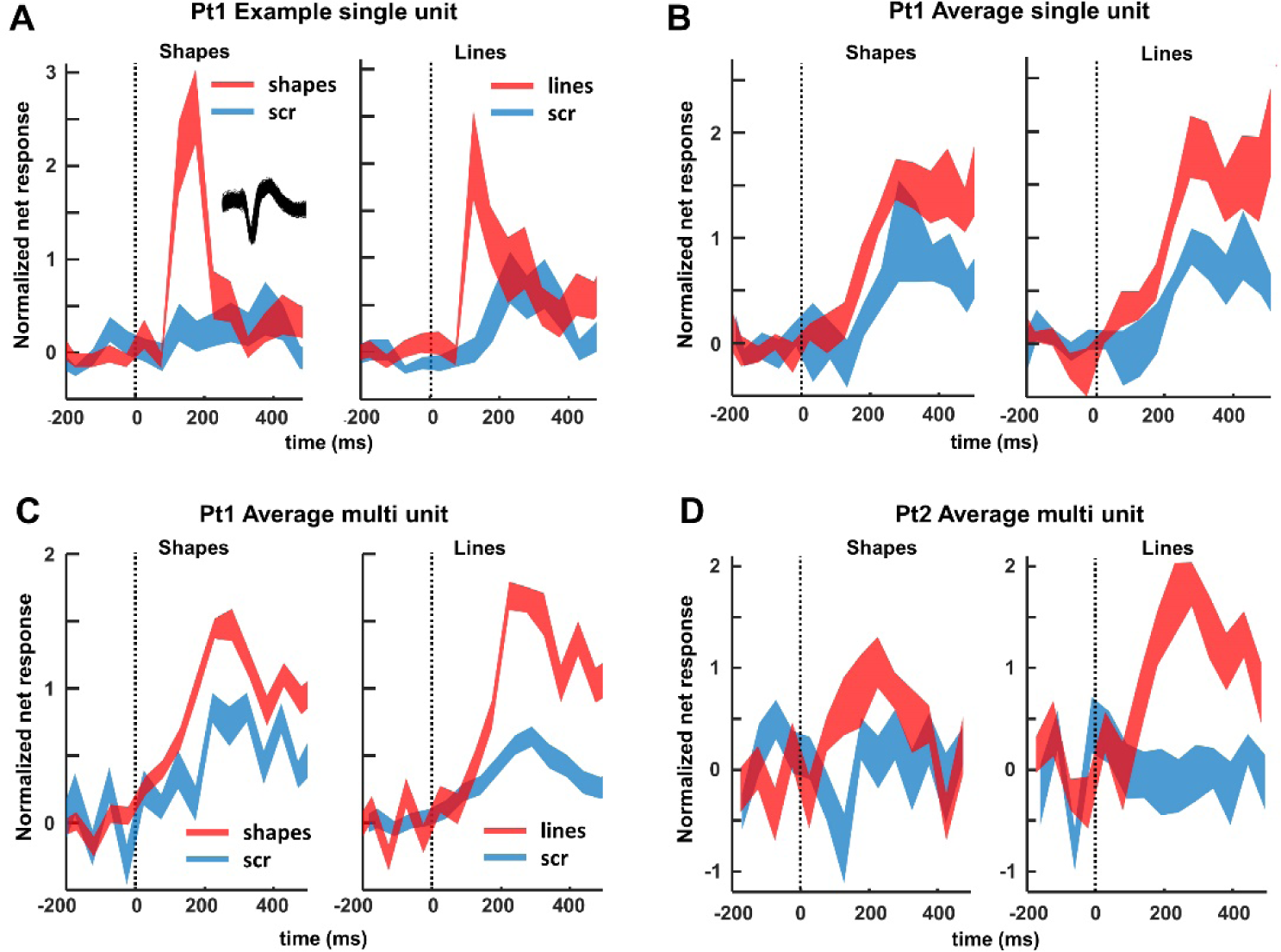
Single- and multiunit responses to images of objects and scrambled controls. A. Example neuron. Average response to intact (red) and scrambled (blue) shapes (left) and line stimuli (right). The inset illustrates the spike waveform. B. Average single-unit responses across all visually responsive channels in patient 1. C and D. Average multiunit responses to intact and scrambled stimuli in patient 1 and 2, respectively, across all visually responsive channels.

We recorded neuronal responses to images of objects in patient 1 in two separate sessions (number of channels: 174), and detected 42 visually-responsive single units (average normalized net response in Fig 2B). Entirely consistent with the fMRI results, half of these neurons (21/42) responded significantly more strongly to intact than to scrambled images of objects (permutation test non-scrambled versus scrambled: p <0.05; median d’ index: 0.35), and no single unit showed a significant preference for scrambled controls. The median response latency of these 21 selective neurons (calculated on the responses to intact shapes and outlines) was 150 ms, whereas the fastest neurons (percentile 10) started to respond at 75 ms after stimulus onset. However, the median selectivity latency (i.e. the first bin with significant response differences between intact and scrambled images) was much higher (225 ms), and no neuron started to discriminate between intact and scrambled stimuli before 75 ms. We obtained highly similar results for MUA in both patients (Figure 2C and D). A large number (46 out of 83) of the visually responsive channels preferred intact over scrambled shapes (41 out of 71 for pt1, 5 out of 12 for patient 2; permutation test: p<0.05; median d’: Pt1: 0.45, Pt2: 0.28). Not surprisingly, the average multi-unit response of all visually-responsive channels was greater for non-scrambled stimuli than for scrambled controls (p<0.001, permutation test, Fig 2C and D).

When looking at the LFP signal that was recorded together with the spiking activity, virtually all channels responded significantly to visual stimulation (80-120Hz, or high-gamma power intact shapes versus pre-stimulus baseline; permutation test, p <0.05; pt1: n = 164/174 or 94%; pt2: n = 269/285 or 94%). On average, we observed a broad-band response after stimulus onset in all four conditions (permutation test, p < 0.01, corrected for multiple comparisons, Figure 3), in which the LFP response to intact shapes and outlines was significantly stronger than to scrambled controls, both at the level of the average high gamma power and on the great majority of the individual channels (permutation test, p < 0.001; pt1: 109 /174 (63%) individual channels; pt2: 242/285 (85%) individual channels). As expected, the lower frequency bands discriminated less reliably between intact and scrambled shapes. Thus, SUA, MUA and LFP data clearly demonstrate that neurons in human LOC are more responsive to intact shapes than to scrambled shapes, confirming and validating the results of the fMRI localizer.

**Figure 3.**
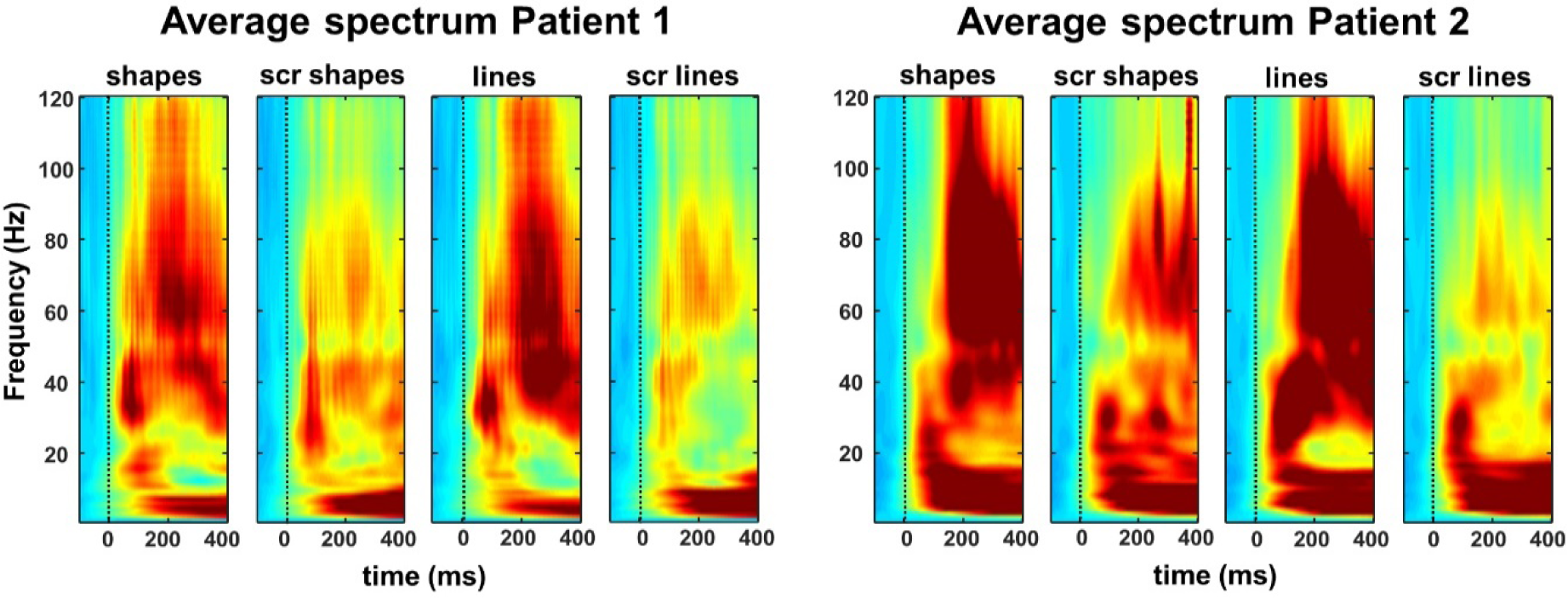
Time-frequency plots of the average local field potentials (LFPs) recorded in the two patients, indicating that the high-gamma (80-120 Hz) power is significantly stronger for intact shapes and outlines than for their scrambled controls.

### Shape selectivity and spatial clustering

MUA recording sites were not only sensitive to image scrambling, but could also be selective for individual shapes (one-way ANOVA p < 0.05 in 8/46 recording sites). To quantify and visualize this shape selectivity, we ranked the intact shapes based on the MUA responses (for all 46 channels with significant selectivity for image scrambling, 6 for pt1, 40 for pt2), and calculated the average MUA response to the ranked intact shapes and to the corresponding scrambled shape images (Fig. 4A and B). Despite the fact that we did not search for selective neurons, shape selectivity was nonetheless robust, since the half-maximum response was measured for rank 17 (patient 1) and 14 (patient 2), and the least-preferred shapes even evoked inhibitory responses. No significant tuning was present for the corresponding scrambled controls (see supplementary Table S1 for linear regression slopes). The results were highly similar for the single-unit responses and for the line stimuli and their scrambled controls (supplementary Figs S1, and S2 and Table S1).

**Figure 4.**
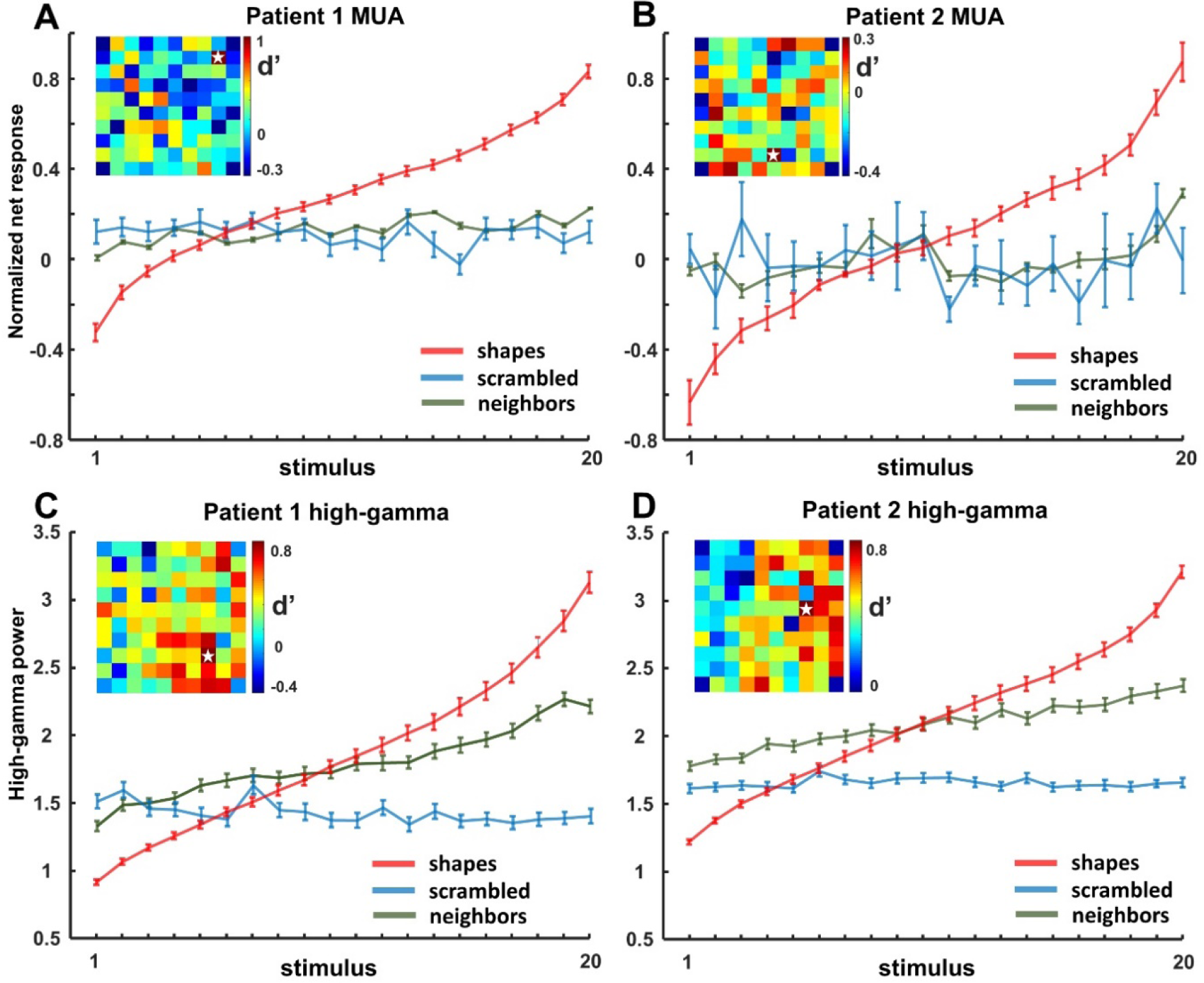
Ranking of shapes for multi-unit activity (MUA) and high-gamma LFP for each patient. The same ranking is applied for the neighboring channels and the corresponding scrambled control stimuli.

The high-gamma responses to intact shapes were equally selective (Figure 4C and D; one-way ANOVA with factor *stimulus number*, for shapes: p < 0.05 in 15/109 for pt1; 72/242 channels for pt2). The most effective intact shape elicited 3 to 4 times more high-gamma activity than the least-preferred shape (half-maximum response for rank 15 in both patients), and no significant tuning was present for the corresponding scrambled controls.

Many MUA and LFP sites were sensitive to image scrambling, but the degree of sensitivity differed markedly on neighboring channels spaced a mere 400 microns apart. This unexpected spatial specificity in the MUA and high-gamma responses to image scrambling became evident in the d’ indices mapped on the spatial layout of the array (compare for example the d’ of a highly selective channel indicated with a star to its neighboring channels, Fig.4A-D, insets). To quantify this spatial specificity to image scrambling across the array, we calculated a 2-way ANOVA for the evoked high-gamma responses for each of the 109 (pt1) or 240 (pt2) selective channels with factors *scrambling* [*scrambled* vs *non-scrambled*] and *position* [neighboring position]. In the large majority of the channels, the main effect of *position* (pt1: N=108/109; pt2: N=235/240) or the interaction between the factors *scrambling* and *position* (pt1: N = 57/109; pt2: N=205/240) were significant (p < 0.05, highly similar results were found for MUA).

We also observed a high degree of spatial clustering for shape preference across the array, both at the level of MUA and at the level of high-gamma responses. For each responsive channel (the center electrode), we calculated the average responses to the intact shapes on all its neighboring channels based on the shape ranking of the center electrode (Fig. 4, green lines). The shape preference differed markedly between each center electrode and its neighbors (Fig. 4 A-D, see Table S1), indicating that the shape preference of human LOC neurons is clustered on a submillimeter scale, similar to the monkey ITC (Fujita et al., 1992, Yamane et al., 2006).

### Receptive fields

A fundamental characteristic of visual neurons is their receptive field (RF). To map the RF of neurons in human visual cortex, we presented an intact shape at 25 positions on the screen covering a 30 by 50 deg area in both hemifields. The four example neurons recorded in patient 1 (Figure 5A) clearly demonstrate that the RFs were relatively large (average surface area 473 deg^2^) and covered both the ipsi- and the contralateral hemifield. Out of 46 visually responsive single neurons (stimulus vs baseline, p <0.05, permutation test), 24 (52%) responded maximally in the contralateral hemifield, 13 (28%) in the ipsilateral hemifield, and 9 neurons responded maximally at the midline (3 of which were at the fovea). Six neurons showed bilateral responses (i.e. > 50 % of the maximal response). The average RF (rightmost panel in Figure 5A) at the single-neuron level included the fovea and the ipsilateral hemifield. The average RF profile was similar when determined using the high gamma responses (Figure 5B): in both patients, the high gamma RF contained the center of the visual field and visual responses (at > 50% of the maximum response) were present both contra- and ipsilaterally. Thus, the average RF in this part of the human LOC was consistently large and bilateral.

**Figure 5.**
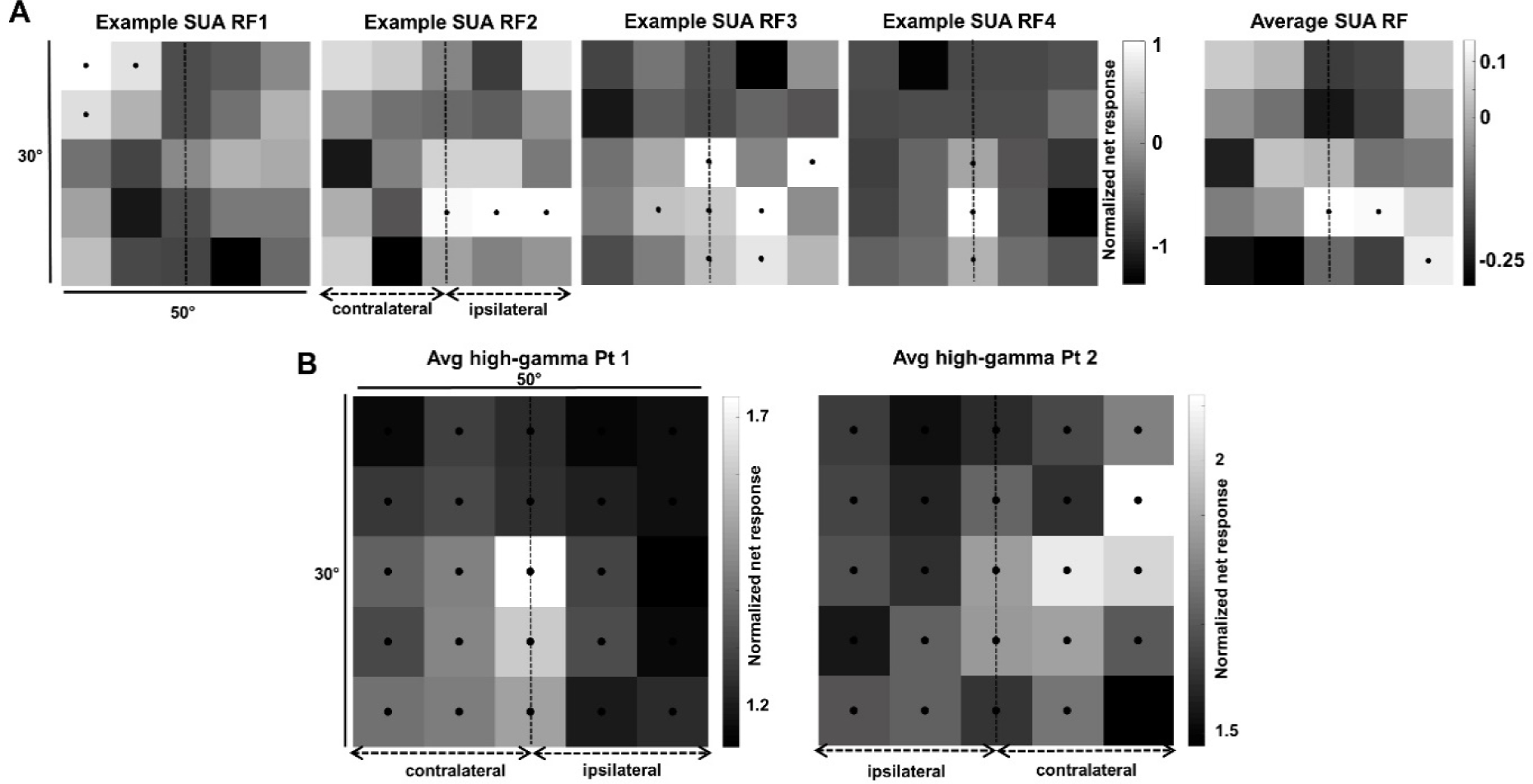
Receptive field mapping. A. Single-unit data. The first four panels show the RFs of four example neurons. All responses are normalized to the maximum visual response. Black dots indicate responses higher than 50% of the maximum response (per channel). Rightmost panel: average receptive field for all visually responsive channels. B. Average RF at the level of high-gamma responses for patient 1 (left) and patient 2 (right).

### Three-dimensional structure selectivity

The previous results have addressed only neural responses to 2D shapes, but neurons in the macaque ITC are also selective for 3D stimuli (Janssen et al., 2000b, Yamane et al., 2008) and several human fMRI studies have suggested that the LOC is sensitive to binocular disparity (Welchman et al., 2005, Georgieva et al., 2008, Moore and Engel, 2001). To investigate the selectivity of LOC neurons for stereo stimuli, we ran a stereo-localizer fMRI experiment, in which blocks of stereo stimuli (curved and flat surfaces at different disparities) alternated with blocks of control stimuli (the monocular images presented without disparity (Durand et al., 2007, Van Dromme et al., 2015)). Figure 6B shows the T-values for the contrast [*stereo*] – [*control*], plotted on the anatomical MRI and CT scans of both patients with the array inserted (p<0.05, FWE corrected). The fMRI results demonstrate that the microelectrode arrays, indicated by the white crosshair, were indeed implanted close to the hotspot of the disparity-related fMRI activations in LOC.

**Figure 6.**
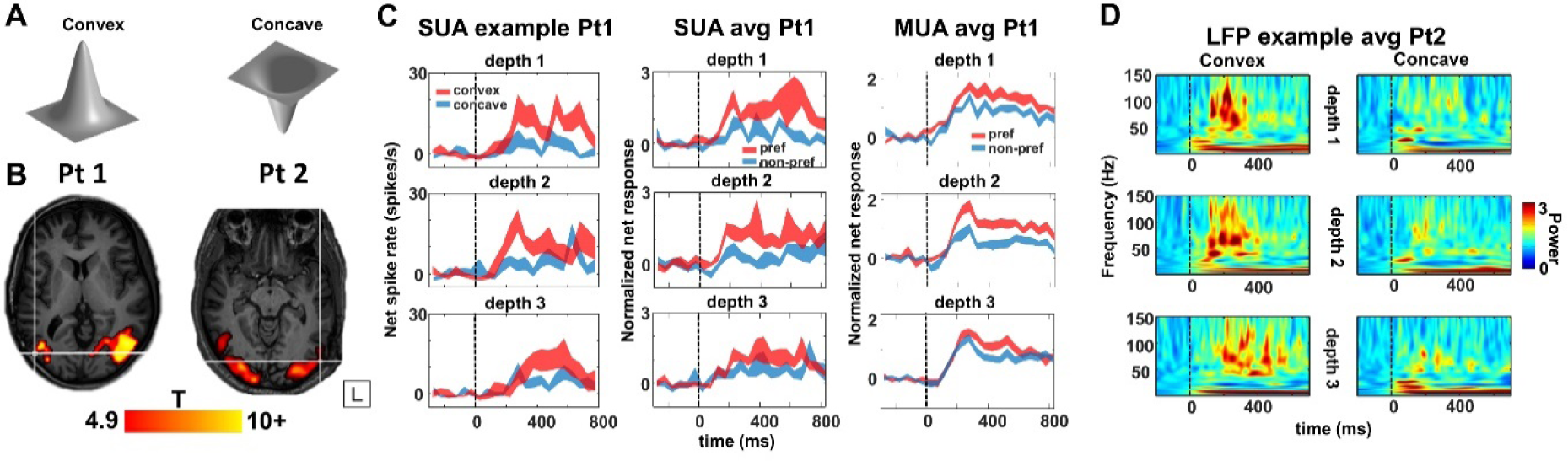
Stereo experiment. A. Stimuli. B. T-values for main effects of stereo, contrast [curved stereo + flat stereo] – [curved control + flat control], plotted on T1 weighted image. P<0.05, FWE-corrected for multiple comparisons. Crosshair indicates the position of the Utah-array. C. Example single neuron (left column), average single-unit responses (middle column) and average multi-unit responses (right column) to preferred (red) and nonpreferred (blue) curved surfaces at three positions in depth (upper row: near, middle row: center and bottom row: far). D. Time-frequency power spectra, for example channel in patient 1, for convex and concave stimulus presentations at three positions in depth. This site is selective for convex shapes across different positions in depth.

We recorded neural activity in LOC during the presentation of stereo stimuli (2 recording sessions) while the patients were categorizing concave and convex surfaces (Fig 6A) at different positions in depth (SUA on 52 channels in Pt 1). The example neuron in Figure 6C (left panel) preferred convex over concave shapes at all three positions in depth (average d’ = 0.73), indicating 3D-structure (i.e. higher-order disparity) selectivity. Notice that this neuron did not start to respond until 100 ms after stimulus onset, and reached its peak activity only after 250 ms. In total, we recorded 39 visually responsive single neurons in this test, 16 (41%) of which showed 3D-structure selectivity (i.e. a main effect of *stereo* and/or a significant interaction between *stereo* and *position in depth* with no reversal in selectivity (Verhoef et al., 2010)). For these 16 selective neurons, we plotted the average net responses to the preferred and non-preferred 3D surfaces at each position in depth (Fig 6C middle panel). This population of LOC neurons preserved its selectivity at every position in depth, as did the MUA (N = 21 sites, right panel in Fig 6C). Similar to the example neuron in Figure 6C, the population (SUA and MUA combined) response latency (125 ms) and the latency of the 3D-structure selectivity (275 ms) were relatively long compared to previously-reported data obtained in the monkey ITC (Verhoef et al., 2012, Janssen et al., 2000b). Moreover, both patients showed significant 3D-structure-selective high gamma responses (15% of visually responsive channels in patient 1, 20% in patient 2, example channel in Figure 6D, average high-gamma of 3D-structure-selective sites is shown for both patients in Figure S3).

Not unlike the selectivity for image scrambling and for individual shapes, the high-gamma 3D-structure preference was highly localized on individual electrodes, since recording sites with a high d’ (convex versus concave) were frequently located next to recording sites with a very low d’ (Supplementary Fig. S4). A 2-way ANOVA with factors *neighboring channel* and *stereo* (convex or concave) indicated a significant clustering of the 3D-structure preference in the large majority of selective channels (94 and 97% of stereo-selective neurons with a main effect of *channel* in patients 1 and 2, respectively, and with 2 and 5 channels, respectively, showing an interaction between channel and stereo in these two patients). Since the high-gamma response correlates with population spiking activity (Liu and Newsome, 2006, Premereur et al., 2012), these results indicate clustering of the 3D-structure preference in human LOC, consistent with previous findings in monkey ITC (Verhoef et al., 2012).

## Discussion

We present the first report of intracortical recordings in SUA, MUA and LFP activity in human LOC using microelectrode arrays. Our 96-electrode array with an interelectrode spacing of 0.4 mm allowed extensive neuronal recordings in human visual cortex with an unprecedented spatiotemporal resolution. Our experiments confirm the robust sensitivity of LOC neurons to image scrambling, as predicted by fMRI, reveal significant 2D-shape and 3D-structure selectivities at the level of SUA, MUA and high-gamma responses, and provide the first RF maps of individual LOC neurons. Moreover, our data furnish new and crucial evidence concerning the microarchitecture of LOC, in that the shape preference differed drastically between neighboring electrodes spaced a mere 400 microns apart.

Our approach using a microelectrode array has several advantages compared to previous electrophysiological studies in humans. Subdural grids with contact points measuring several millimeters (Yoshor et al., 2007) sample neural activity over a wide area, whereas we used intracortical microelectrodes with sharp tips spaced just 400 microns apart, so that a single row of 10 electrodes occupied a stretch of cortex measuring only 3.6 mm. Furthermore, the microelectrode array allowed us to simultaneously record neural activity on 96 microelectrodes, compared to recordings on only two microelectrodes in (Self et al., 2016). Finally, the spatial arrangement of the microelectrode array (10 by 10 electrodes) also allowed us to investigate the microarchitecture of human visual cortex at the scale of cortical columns. It should also be noted that our recording sites were located near the entry point of one of the depth electrodes and were, in retrospect, not part of the epileptogenic zone.

Ever since the original publication by (Kourtzi and Kanwisher, 2000), numerous studies have used the LOC localizer [intact shapes – scrambled shapes] to identify shape-sensitive regions in human visual cortex. However, the actual underlying neural selectivity has never been revealed. An extensive body of work has employed visual adaptation, observed via fMRI activation, as an indirect measurement of neuronal shape selectivity in humans (e.g. (Grill-Spector and Malach, 2001)), but the interpretation of these effects and their relation to neural selectivity at the single-cell level remain controversial (Sawamura et al., 2006). Here, we not only confirmed the strong effect of image scrambling on SUA, MUA and high-gamma responses in LOC, but we also revealed significant response differences for intact shapes, i.e. shape selectivity, at the level of single neurons, as previously shown in the macaque ITC (Logothetis and Sheinberg, 1996, Tanaka, 1996). A bilateral lesion of LOC produces a profound deficit in shape recognition (Goodale et al., 1991, James et al., 2003, Westwood and Goodale, 2011), similar to ITC lesions in monkeys (Cowey and Gross, 1970, Britten et al., 1992, Dean, 1976, Gross, 1994, Dean, 1979), and Transcranial Magnetic Stimulation over LO also impairs shape discrimination (Chouinard et al., 2017). Our data support the notion that these deficits arise from a loss of shape-selective neurons in LOC.

A substantial fraction of our MUA and LFP recording sites showed significant shape tuning – possibly as strong as in the monkey ITC (De Baene and Vogels, 2010) – indicating that shape preference may be localized within the LOC. In addition, the fixed arrangement of the microelectrodes, spaced 400 microns apart, allowed assessing the spatial organization of sensitivity to image scrambling and of the shape selectivity at a high spatial resolution. Our observation that the shape preference changed markedly over the extent of 400 microns is in line with previous studies in the macaque ITC demonstrating considerable clustering for shape features. Fujita et al ((Fujita et al., 1992)) showed that ITC neurons with similar shape selectivities are organized vertically across the cortical thickness, and (Tsunoda et al., 2001) used intrinsic optical imaging to highlight patches of activation elicited by specific object images spaced 0.4 to 0.8 mm apart. Thus, our results are highly consistent with previous studies in the ITC of macaque monkeys.

The spatially-restricted nature of high-gamma responses we measured in LOC is consistent with previous studies in visual cortex indicating that the higher frequency bands of the LFP signal correspond with MUA (Goense and Logothetis, 2008, Liu and Newsome, 2006, Logothetis et al., 2001, Premereur et al., 2012) and originate from a small region of cortex measuring a few hundred microns in extent (Katzner et al., 2009). However, studies in auditory cortex reported considerable volume conduction – even in the high-gamma band of the LFP signal – over several millimeters of cortex (Kajikawa and Schroeder, 2011). Although our data do not allow us to fully resolve this controversy, it should be noted that our spatially-selective recordings were obtained during an active fixation task, where no influence of anesthetics was possible, in contrast to the Kajikawa and Schroeder study. It should also be noted that although we recorded SUA in one patient, we were able to confirm our findings with MUA and high-gamma responses in both patients.

Our data also shed light on the RF properties of LOC neurons. Although we did not obtain sufficient data for an exhaustive RF description, a few conclusions seem to be warranted. The average RF size in LOC was large (473 deg^2^), and a considerable fraction of LOC neurons responded to stimuli presented in the ipsilateral hemifield. A previous study in human early visual cortex reported that the RF measured with high-gamma responses is even more restricted than that measured with MUA (Self et al., 2016). Thus, our observation that LOC sites frequently exhibit bilateral SUA, MUA and high-gamma responses is interesting in view of the comparison between the human LOC and monkey ITC areas (see below). Future studies will have to investigate the RF profile of LOC neurons in more detail.

To our knowledge, we also provide the first evidence for 3D-structure selectivity defined by binocular disparity in human visual cortex. A number of recording sites showed differential responses to convex and concave surfaces (composed of the same monocular images), across different positions in depth, indicative of higher-order disparity or 3D-structure selectivity. Similar to the selectivity for image scrambling and that for individual shapes, the 3D-structure preference was highly localized on individual electrodes, since recording sites with strong selectivity were frequently located next to recording sites with a very low selectivity. A large number of studies (Janssen et al., 2001, Janssen et al., 2003, Janssen et al., 1999, Janssen et al., 2000b, Janssen et al., 2000a, Yamane et al., 2008) have investigated 3D-structure selectivity in the macaque anterior ITC (area TE). More recently, Verhoef et al. showed clustering of the MUA selectivity for 3D structure in area TE using identical stimuli (Verhoef et al., 2012), and microstimulation of these clusters could predictably alter the perceptual report of the animal in a 3D-structure categorization task. Hence, the 3D-structure selectivity we observe here has also been described in the macaque ITC. Moreover, patient DF (Read et al., 2010), who suffered bilateral damage to LOC, was impaired in using the relative disparity between features at different locations, although 3D-structure categorization was not tested.

One major, outstanding question relates to the possible homologies between the different subparts of human LOC and monkey ITC areas TEO and TE (Orban et al., 2014). Although a single study cannot resolve this homology question, several observations we made are highly relevant in this respect. Shape-selective responses can be observed in TEO and in TE. However, the robust 3D-structure selectivity we observed was previously described in the more anterior part of area TE in macaques (Janssen et al., 2000b), but is virtually absent in macaque area TEO (Alizadeh et al., 2018). Moreover, the large and frequently bilateral RFs we observed in human LOC are more consistent with TE than with TEO (Hikosaka, 1999). Detailed mapping of receptive fields (Boussaoud et al., 1991, Kobatake and Tanaka, 1994) and stimulus reduction during single-unit recordings (Tanaka et al., 1991) will clarify in greater detail the relationship between this part of the LOC and the monkey ITC areas.

Overall, the similarities between the human LOC and the (more anterior part of) monkey ITC were very apparent. In both species, neurons are sensitive to image scrambling (Vogels, 1999) and shape-selective; shape preference is clustered; the RFs are large and include the fovea (Op de Beeck et al., 2001), and neurons preserve their 3D structure preferences across position in depth (Janssen et al., 2000b). The only possible discrepancy between our results in the human LOC and previous studies in monkey ITC may reside in the latencies of the neuronal response. When tested with images of objects, the MUA became selective only after 125 ms, and the response and selectivity latency for 3D stimuli equaled 125 and 225 ms, respectively. In contrast, macaque ITC neurons can signal shape differences starting at 70-80 ms after stimulus onset and at 80-100 ms for 3D stimuli (Janssen et al., 2000b)). However, our results are in line with previous studies in the human medial temporal lobe – which is downstream from LOC – reporting latencies of 300 ms (Quiroga et al., 2005, Kreiman et al., 2000b). However, neuronal latencies are highly influenced by the number of stimulus repetitions, therefore a higher number of selective recording sites and a higher number of trials may have yielded shorter latencies in our experiments. More detailed measurements in the two species in areas where the homology is clearly established (e.g. V1) are undoubtedly necessary.

## Materials and Methods

Two patients (Pt2, 28 y.o. female; Pt1, 44 y.o. male) with refractory epilepsy underwent invasive intracranial recordings with the use of depth electrodes to delineate the epileptogenic zone. During this procedure microelectrode arrays were implanted in the left (patient 1) and right (patient 2) lateral occipital complex. Informed consent from the patients and local ethical board approval was obtained for this procedure (study protocol S 53126).

### fMRI

#### Stimuli

The stimuli were projected from a liquid crystal display projector (Barco Reality 6400i, 1024 × 768 pixels, 60-Hz refresh rate) onto a translucent screen positioned in the bore of the magnet (57 cm distance). The patients viewed the stimuli through a mirror tilted at 45° and attached to the head coil.

##### LO Localizer

The LO localizer stimuli were images measuring 300 × 300 pixels. We used grayscale images and line drawings of familiar objects (20 images and 20 line drawings), as well as scrambled versions of each set (Fig. 1A) (Kourtzi et al., 2003). The scrambled images were created by dividing the intact images into a 20 × 20 square grid and randomizing the positions of each of the resulting squares. The grid lines were present in both the intact and the scrambled images. The overall size of the stimuli measured 7° in visual angle and each stimulus was presented for 1000 ms.

##### Stereo Localizer

The stimulus set consisted of random-dot stereograms in which the depth was defined by horizontal disparity (dot size 0.08°, dot density 50%, vertical extent 5.5°), and were presented on a gray background. All stimuli were generated using MATLAB (R2010a, MathWorks) and were gamma-corrected. We used a 2 by 2 design with factors *curvature* (curved vs flat) and *disparity* (stereo vs control), as described in (Durand et al., 2007, Joly et al., 2009, Van Dromme et al., 2016, Van Dromme et al., 2015). The stereo-curved condition consisted of three types of smoothly-curved depth profiles (1, 1/2, or 1/4 vertical sinusoidal cycle) together with their antiphase counterparts obtained by interchanging the right and left monocular images (disparity amplitude within the surface: 0.5°). Each of the six depth profiles was combined with one of four different circumference shapes and appeared at two different positions in depth (mean disparity + or − 0.5°), creating a set of 48 curved surfaces. In the stereo-flat condition, flat surfaces (using the same four circumference shapes) were presented at 12 different positions in depth, such that the disparity content (the sum of all disparities) was identical to that in the stereo curved condition. Finally, the control conditions (stereo-control and flat-control) consisted of the presentation of one of the monocular images (either belonging to one of the stereo-curved stimuli or to one of the stereo-flat stimuli) to both eyes simultaneously. Each control condition consisted of exactly the same monocular images as the corresponding stereo condition, hence the binocular input was identical in the stereo conditions and in the control conditions. The overall size of the stimuli measured 5.6° in visual angle and each stimulus was presented for 1000 ms. Dichoptic presentation of the stimuli was achieved by means of red/green filter stereo glasses worn by the patient.

#### Data collection

Scanning was performed on a 3-T MR scanner (Achieva dstream, Philips Medical Systems, Best, The Netherlands) located at the University Hospitals Leuven. Functional images were acquired using gradient-echoplanar imaging with the following parameters: 52 horizontal slices (2 mm slice thickness; 0.2 mm gap; multiband acquisition), repetition time (TR) 2 s, time of echo (TE): 30 ms, flip angle: 90°, 112 * 112 matrix with 2 ×2 mm in-plane resolution, and sensitivity-enhancing (SENSE) reduction factor of 2. The 25 slices of a volume covered the entire brain from the cerebellum to the vertex. A three-dimensional (3D) high-resolution (18181.2) T1-weighted image covering the entire brain was acquired in the beginning of the scanning session and used for anatomical reference (TE/TR 4.6/9.7 ms; inversion time, 900 ms; slice thickness, 1.2 mm; 256 * 256 matrix; 182 coronal slices; SENSE reduction factor 2.5). The single scanning session lasted 60 min.

##### CT

A computed tomography (CT) scan (Siemens, 1mm slice thickness, 120kV, Dose length product of 819mGy.cm) was performed two hours after electrode placement to verify the location of the microelectrode array.

##### LO Localizer

Stimuli (shapes, line stimuli, scrambled shapes, scrambled line stimuli, fixation only, Fig. 1A) were presented in blocks of 24 s except for the fixation condition (20 s), each block was repeated 4 times in a run, creating runs of 464 s. Individual stimuli were presented for 1000 ms (ISI=0; fixation time: 200 ms).

##### Stereo Localizer

Stimuli (curved stereo, flat stereo, curved control, flat control, fixation only, Fig. 1B) were presented in blocks of 24 s, and each block was repeated 4 times in a run, creating runs of 480 s. Individual stimuli were presented for 1000 ms (interstimulus interval = 0 ms; fixation time = 200 ms). 12 functional volumes were acquired for every block (or condition, each 24 s long) and these were embedded in a time series of 222 volumes (444 s).

#### Data analysis

Data analysis was performed using the SPM12 software package (Wellcome Department of Cognitive Neurology, London, UK) running under MATLAB (The Mathworks, Natick, MA). The preprocessing steps involved *1*) realignment of the images, *2*) coregistration of the anatomical image and the mean functional image. Before further analysis, the functional data were smoothed with an isotropic Gaussian kernel of 5 mm. To determine the exact location of the Utah array, the CT scan was coregistered with the anatomical image using SPM12 software.

##### LO Localizer

To localize areas responding more strongly to the presentation of objects versus scrambled controls, we calculated the contrast [shapes + outlines] – [scrambled shapes +scrambled outlines], at p < 0.05, FWE corrected.

##### Stereo

To identify regions sensitive to binocular disparity, we calculated the main effect of *stereo*: [curved stereo + flat stereo] – [curved control + flat control], at p < 0.05, FWE corrected.

### Electrophysiology

We implanted a 96-channel microelectrode array with 1.5 mm electrode length in patient 1 and with 1 mm electrode length in patient 2; electrode spacing measured 400 microns (4 × 4 mm; Blackrock Microsystems, UT, USA). The array was implanted through a burr hole over the occipitotemporal cortex, used for depth electrode placement, according to the manufacturer’s protocol with a pressurized inserter wand. These microelectrodes were used clinically for advanced epilepsy monitoring (study protocol S 53126).

#### Stimuli

All stimuli were presented by means of a custom-made stereoscope. Images from two LCD monitors were presented to the two eyes with the use of customized mirrors at a viewing distance of 56 cm (1 pixel = 0.028°). Continuous eye-movement tracking (left eye, 120Hz; ISCAN, MA, USA), ensuring fixation in an electronically defined window (3*3 degrees), was performed throughout the experiment. Trials in which the patients did not maintain fixation were aborted.

##### LO Localizer

The same stimulus set as in the fMRI experiment was used. The the stimuli presented in the stereoscope were 8.5 deg in size. After a brief period of fixation (200 ms), the stimulus was presented for 500 ms, followed by an interstimulus interval of 100 ms. In the LO localizer, no disparity was present in the stimuli.

##### Receptive field mapping

To map the RF, a single non-scrambled shape (8.5 deg) was presented at 25 different positions in the visual field, covering 50 degrees horizontally and 30 degrees vertically, during passive fixation.

##### Stereo test

We presented concave and convex surfaces at three different positions in depth (near, at the fixation plane, and far, Fig. 1C) at the fixation point while monitoring the position of the left eye. To avoid monocular depth cues, the disparity (disparity amplitude: 0.25 deg) varied only along the surface of the shape, while the circumference of the shape was kept at a constant disparity (+0.25 deg, 0 deg or −0.25 deg disparity), as in (Verhoef et al., 2010, Verhoef et al., 2012). The patients had to categorize the 3D structure of the stimulus (concave or convex, 100% disparity coherence) independently of the position in depth by means of a button press after stimulus offset (1000 ms of stimulus presentation time), as in (Verhoef et al., 2010). An auditory tone provided feedback after every successfully completed trial. Both patients performed at more than 90% correct.

#### Data Collection

Data were collected using a digital headstage (Blackrock Microsystems, UT, USA) connected to a 128-channel neural signal processor (Blackrock Microsystems, UT, USA). For LFP recordings, the signal was filtered with a digital low-pass filter of 125 Hz, and LFP signals were recorded continuously (sampling frequency: 1000 Hz). Single- and multi-unit signals were high-pass filtered (750 Hz). A multi-unit detection trigger was set at a level of 95% of the signal’s noise. All spike sorting was performed offline (Offline Sorter 4, Plexon, TX, USA).

#### Data analysis

Data analysis was performed using custom-written Matlab (the MathWorks, MA, USA) software.

##### Spike rate analysis

For every channel, we calculated the net spike rate by subtracting the average baseline activity from the spike rate. Spike rate was further normalized by dividing the net spike rates by the average spike rate for the best condition (50-300 ms after stimulus onset) for each channel. Statistics were performed using permutation tests, where real data were randomly distributed over all the different conditions 1000 times. The differences between two conditions were calculated for every permutation, and compared with the actual difference between conditions. The latency of the spiking activity for visually-responsive channels was defined as the first of two consecutive 50 ms bins with a spike rate higher than the average baseline plus 2 standard errors. The selectivity latency was defined as the first of two consecutive 50 ms bins with a spike rate for the preferred condition higher than the average spike rate for the non-preferred condition plus 2 standard errors.

##### LFP analysis

For every trial, the time-frequency power spectrum was calculated using Morlet’s wavelet analysis techniques (Tallon-Baudry and Bertrand, 1999), with a spectro-temporal resolution equal to 7, after filtering with a 50 Hz notch filter (FieldTrip Toolbox, Donders Institute, The Netherlands (Oostenveld et al., 2011)). This method provides a better compromise between time and frequency resolution compared to methods using Fourier transforms (Sinkkonen et al., 1995, Tallon-Baudry et al., 1997). To remove any filter artifacts at the beginning and end of the trial, the first and last 100 ms of each trial were discarded. Power was normalized per trial by dividing the power trace per frequency by the average power for this frequency in the 100 ms interval before stimulus onset. To exclude trials containing possible artifacts in the LFP recordings, maximum values of the continuous LFP signal were determined and trials with maximum values above the 95^th^ percentile were removed. Furthermore, the data set was split in two, and all population analyses were repeated for both halves of the data independently, to check for consistency. We analyzed the LFP power in the high frequency bands (high-gamma): 80-120 Hz. Lowest frequencies had to be excluded from our analyses, as our trials were maximally 1 s long. All statistics on LFP data were obtained using permutation tests as described for spiking activity. The latency of the LFP response per frequency band was defined as the first of five consecutive timestamps (in ms) in which the average power minus 2 standard errors was higher than 1 (= average power of the normalized baseline). The LFP-latency for selectivity between conditions was defined as the first of two consecutive samples in which the average power for condition *A* minus 2 standard errors was higher than the average power for condition *B*.

##### D-prime

d’ statistics were calculated as:

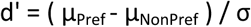

Here μ_Pref_ and μ_NonPref_ denote the mean responses to the preferred and the non-preferred condition (e.g. non-scrambled versus scrambled), respectively, and

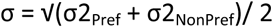

is the pooled variance of the two response distributions. This measure differs from those used in previous studies (Frien and Eckhorn, 2000, Frien et al., 2000, Liu and Newsome, 2006) in that it explicitly takes into account the trial-by-trial variability of the response (Siegel and Konig, 2003).

To estimate the spatial extent of selectivity observed in the MUA and high-gamma band over the array, we determined for each visually responsive channel its immediate neighbors (i.e. either 8 channels, for recording channels which were not located on the edge of the array, or 5 channels for edge electrodes). We calculated a two-way ANOVA with factors *scrambling* (scrambled vs non-scrambled) and *position* for each channel individually. Significance was tested using a p-value of 0.05.

##### Ranking

*T*o investigate the MUA and high-gamma responses to individual stimuli, we applied a ranking technique in which individual non-scrambled stimuli were ranked based on the electrode’s average spiking activity and high-gamma power evoked by the stimuli, then the same ranking was applied to the individual scrambled control stimuli. To investigate differences between rankings, a linear regression was performed, and a 95% confidence interval was used to determine significant differences between regression coefficients or intercepts. Finally, the same ranking technique was used to investigate the spatial specificity of the shape selectivity: we ranked the non-scrambled stimuli for each electrode based on the spiking activity and high-gamma responses, and this same ranking was then applied to the responses of all neighboring channels separately. We then averaged the spike rate and gamma responses for the ranked data of the neighboring channels to determine whether the shape preference was preserved at neighboring channels. Differences in ranking were investigated using a linear fit as described above.

##### Receptive fields

The average single-unit activity and high gamma power were calculated during stimulus presentation for each stimulus-position, and filtered with a Gaussian (sigma: 0.5).

To calculate receptive field size, we constructed RF maps by interpolating the neuronal responses between all positions tested across the 50*30 degrees display area, and then calculating the number of pixels in the RF map with a response higher than 50% of the maximum response.

## Author contributions

TD, EP, PJ and TT designed experiment, TD, MU and TT collected data, EP, TD performed data-analysis, TD, EP, PJ, TT, WVP, and JvL wrote the paper

## Supplementary Information

**Figure S1.**
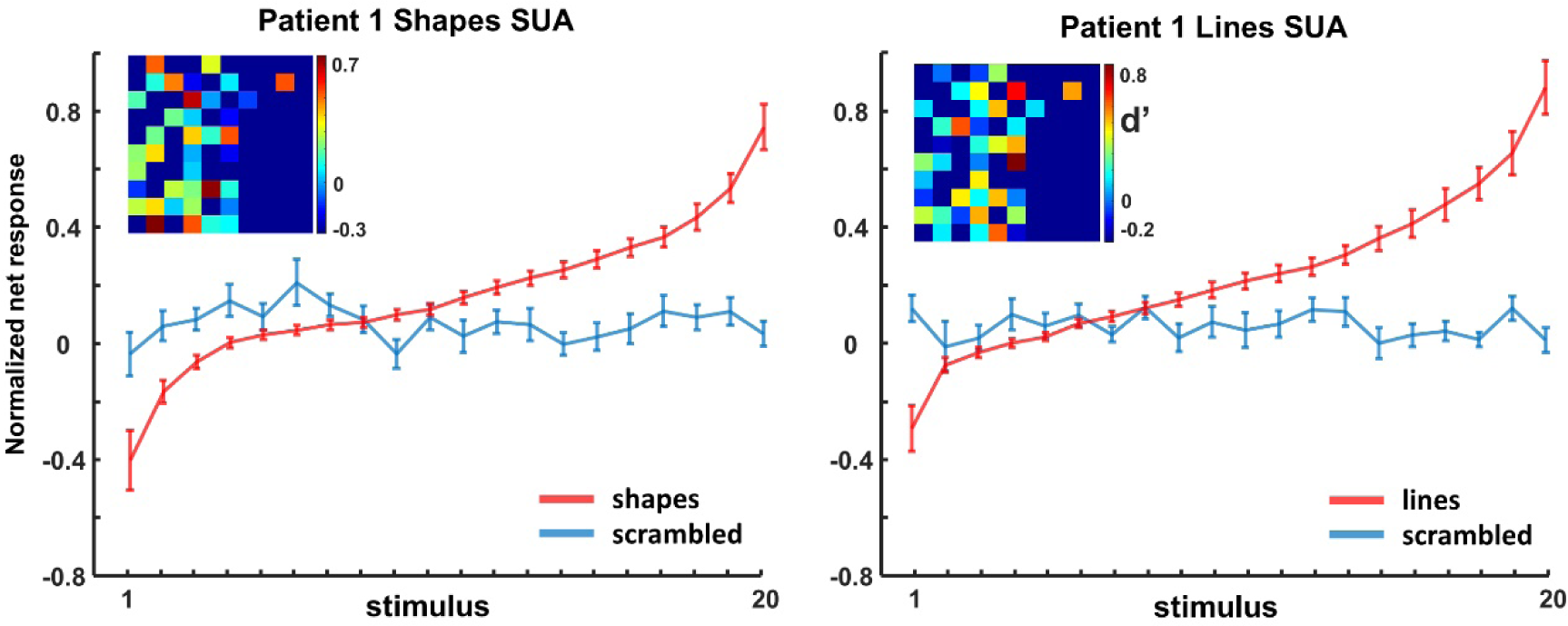
Ranking analyses of shapes and lines for single-unit activity (SUA) in patient 1. The same ranking is applied for the corresponding scrambled control stimuli.

**Figure S2.**
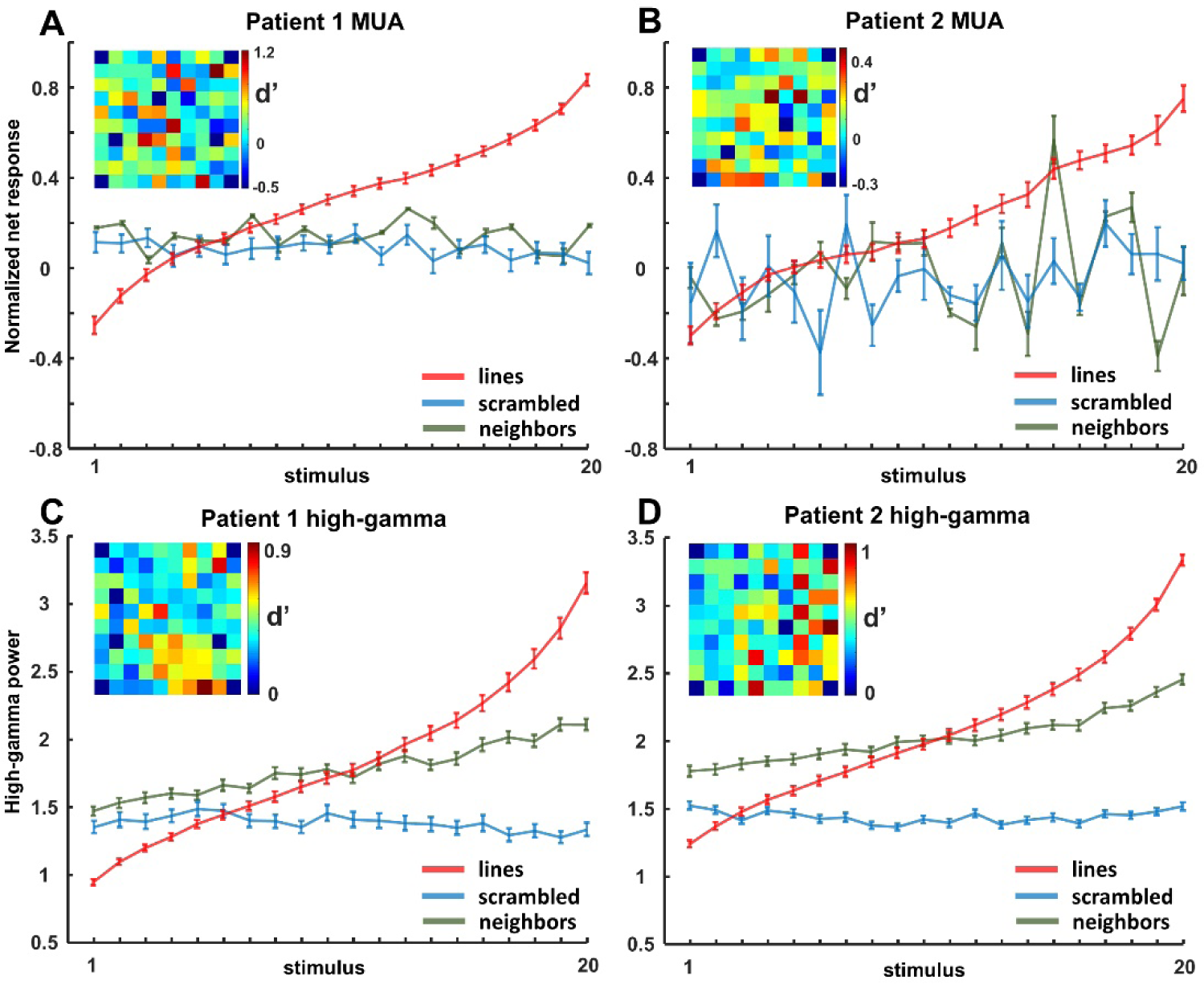
Ranking of lines for multi-unit activity (MUA) and high-gamma LFP for both patients. The same ranking is applied for the neighboring channels and the corresponding scrambled control stimuli.

**Figure S3.**
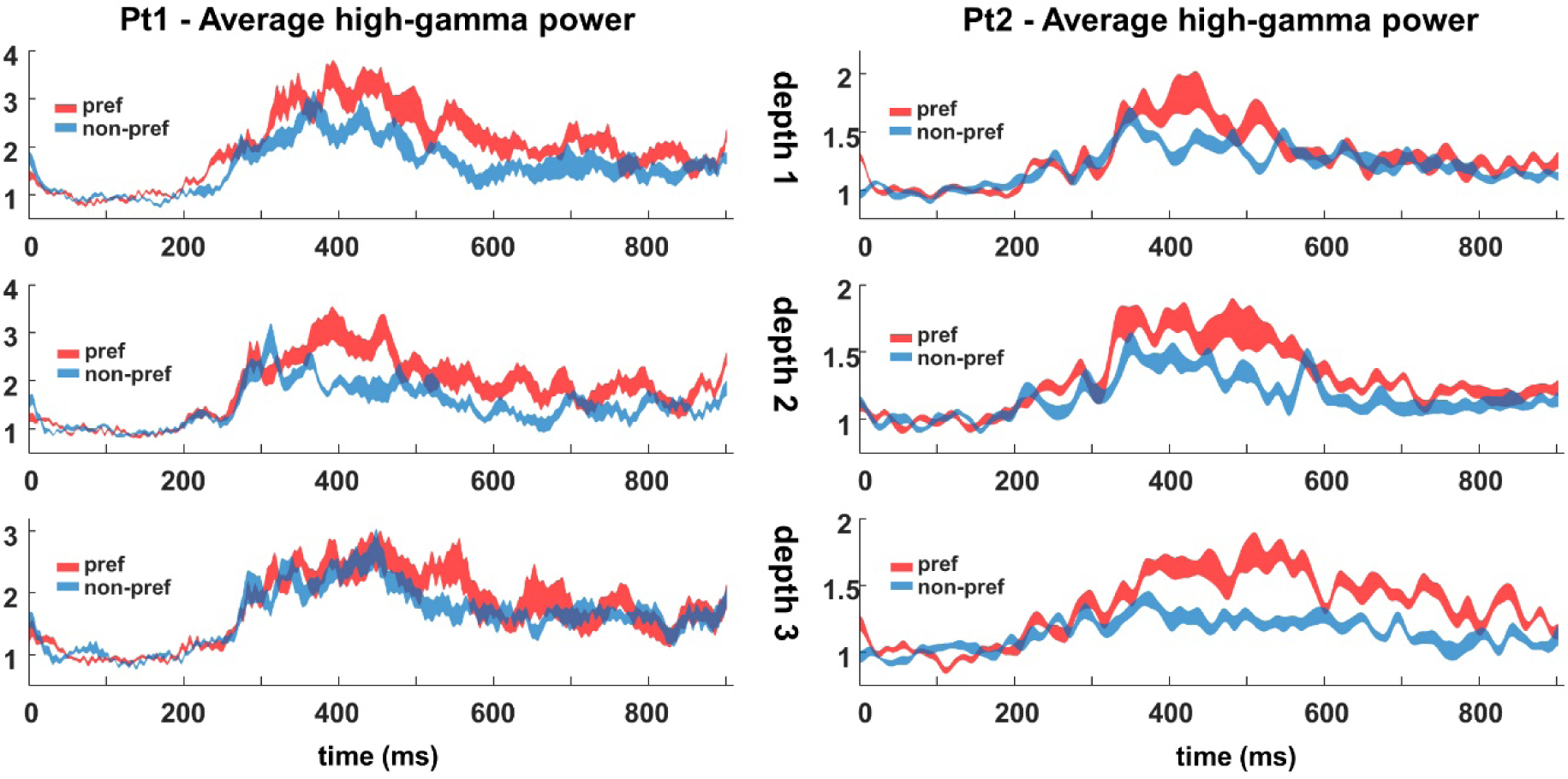
Average high-gamma of 3D-structure-selective sites for each patient. Preferred vs non-preferred shapes.

**Figure S4.**
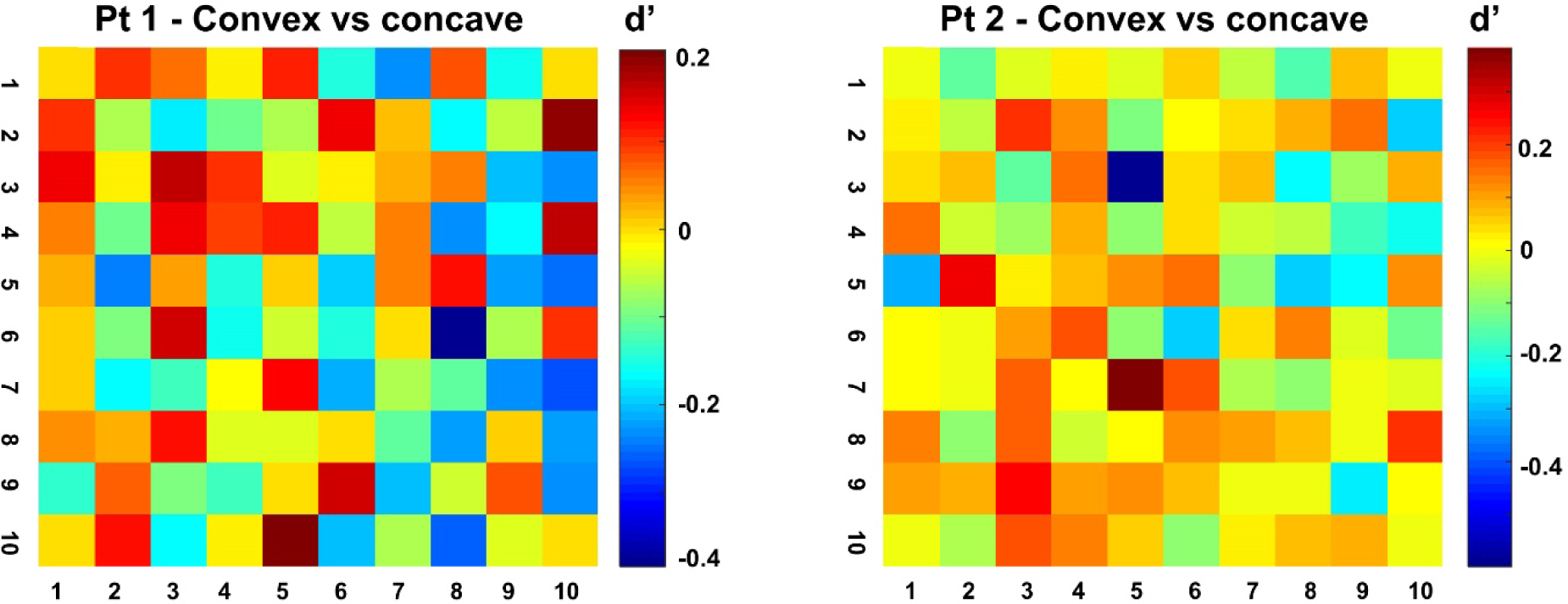
d’ values of high-gamma responses for the two patients across the array. High-gamma 3D-structure preference was highly localized on individual electrodes, since recording sites with a high d’ (convex versus concave) were frequently located next to recording sites with a low d’.

**Table S1.** Slope of regression lines with 95% confidence interval for shapes and lines. The same ranking was applied for the scrambled versions of the stimuli and for their neighboring channels respectively. * Indicates significant ranking difference.

|  | **Patient 1** |  |  | **Patient 2** |  |
| --- | --- | --- | --- | --- | --- |
|  | *SUA* | *MUA* | *LFP* | *MUA* | *LFP* |
| **Shapes** | 0.0404<br>(0.0337, 0.047) | 0.049<br>(0.0448, 0.0531) | 0.1015<br>(0.0932, 0.1097) | 0.0623<br>(0.0561, 0.0685) | 0.0892<br>(0.0834, 0.095) |
| **Applied ranking for scr_shapes** | -0.0008*<br>(-0.0058, 0.0042) | -0.0026*<br>(-0.0064, 0.0011) | -0.0077*<br>(-0.0129, -0.0026) | -0.0005*<br>(-0.0098, 0.0087) | 0.0002*<br>(-0.0027, 0.0031) |
| **Applied ranking for neighbors** | / | 0.0072*<br>(0.0043 - 0.0101) | 0.041*<br>(0.0362 - 0.0458) | 0.0082*<br>(0.001 - 0.0153) | 0.0285*<br>(0.026 - 0.0309) |
| **Lines** | 0.0443<br>(0.038, 0.0507) | 0.0468<br>(0.0431, 0.0505) | 0.0974<br>(0.0867, 0.108) | 0.0469<br>(0.0432, 0.0506) | 0.0919<br>(0.082, 0.1018) |
| **Applied ranking for scr_lines** | -0.0009*<br>(-0.289, -0.0047) | 0.1483*<br>(-0.0028, -0.0056) | -0.0059*<br>(-0.0094, -0.0024) | 0.0073*<br>(-0.0047, 0.0194) | -0.0005*<br>(-0.0043, 0.0034) |
| **Applied ranking for neighbors** | / | -0.0006*<br>(-0.0057 - 0.0045) | 0.0312*<br>(0.0279 - 0.0346) | 0.0007*<br>(-0.0116 - 0.0256) | 0.0305*<br>(0.0262 - 0.0347) |

## Acknowledgments

We thank Stijn Verstraeten, Piet Kayenbergh, Gerrit Meulemans, Marc De Paep, Anaïs Van Hoylandt, Ron Peeters and Evy Cleeren for technical assistance. We thank Astrid Hermans and Sara De Pril for administrative support.

This work was supported by Fonds Wetenschappelijk onderzoek (FWO) and Odysseus grant. T.T. is supported by FWO (senior clinical researcher, FWO 1830717N).

